# SARS-CoV-2 ORF8 is a viral cytokine regulating immune responses

**DOI:** 10.1101/2022.08.01.502275

**Authors:** Masako Kohyama, Tatsuya Suzuki, Wataru Nakai, Chikako Ono, Sumiko Matsuoka, Koichi Iwatani, Yafei Liu, Yusuke Sakai, Atsushi Nakagawa, Keisuke Tomii, Koichiro Ohmura, Masato Okada, Yoshiharu Matsuura, Shiro Ohshima, Yusuke Maeda, Toru Okamoto, Hisashi Arase

## Abstract

Many patients with severe COVID-19 suffer from pneumonia, and thus elucidation of the mechanisms underlying the development of such severe pneumonia is important. The ORF8 protein is a secreted protein of SARS-CoV-2, whose in vivo function is not well understood. Here, we analyzed the function of ORF8 protein by generating ORF8-knockout SARS-CoV-2. We found that the lung inflammation observed in wild-type SARS-CoV-2-infected hamsters was decreased in ORF8-knockout SARS-CoV-2-infected hamsters. Administration of recombinant ORF8 protein to hamsters also induced lymphocyte infiltration into the lungs. Similar pro-inflammatory cytokine production was observed in primary human monocytes treated with recombinant ORF8 protein. Furthermore, we demonstrate that the serum ORF8 protein levels are correlated well with clinical markers of inflammation. These results demonstrated that the ORF8 protein is a viral cytokine of SARS-CoV-2 involved in the in the immune dysregulation observed in COVID-19 patients, and that the ORF8 protein could be a novel therapeutic target in severe COVID-19 patients.

## Introduction

The serious consequences of SARS-CoV-2 infection is the immune dysregulation or inflammatory response that leads to severe pneumonia, which is directly related to the death of patients with COVID-19 (1-4). In addition, prolonged SARS-CoV-2 infection is observed in some COVID-19 patients, suggesting the presence of an immune evasion mechanism (5,6). SARS-CoV-2 is a positive-sense RNA virus, possessing four structural, sixteen non-structural, and six accessory proteins. ORF8 encodes one of the accessory proteins that is conserved among coronaviruses. The SARS-CoV-2 ORF8, but not the SARS-CoV-1 ORF8, contains a N-terminal signaling sequence, suggesting that the SARS-CoV-2 ORF8 might be a newly acquired secretory protein (7,8), and recently it has been reported that SARS-CoV-2 ORF8 is secreted protein (9). Moreover, anti-ORF8 antibodies are detected in COVID-19 patients (10), suggesting that the ORF8 protein is produced upon SARS-CoV-2 infection. Several recent papers suggested the possible function of ORF8 protein; however, as they were based on the transfection of ORF8, the physiological function of ORF8 in SARS-CoV-2 infection has remained unclear (11-14). Here we demonstrated that ORF8 protein is involved in lung inflammation by generating ORF8-knockout SARS-CoV-2 and recombinant ORF8 protein. Moreover, we found that serum ORF8 levels were correlated well with clinical markers of inflammation in COVID-19 patients.

## Materials and Methods

### Human sample

The collection and use of human sera and saliva were approved by Osaka University (2020-10 and 19546), Kobe City Medical Center General Hospital (200924), and Osaka South Hospital (2-28). Written informed consent was obtained from the participants according to the relevant guidelines of the institutional review board. The diagnoses of SARS-CoV-2 were PCR-based. The patients requiring a ventilator were considered as severe patients. All sera from the SARS-CoV-2 patients were treated with 2% CHAPS for 30 min at room temperature to inactivate the remaining virus.

### Cell lines

HEK293T cells (RIKEN Cell Bank), THP-1 (Japanese Collection of Research Bioresources Cell Bank, JCRB0112), TMPRSS2-expressing VeroE6 cells (JCRB1819), and IFNAR1-deficient HEK293-3P6C33 cells in which human ACE2 and TMPRSS2 are induced by doxycycline were cultured in DMEM or RPMI (Nacalai, Japan) supplemented with 10% FBS Japan) and cultured at 37C in 5% CO2. The Expi293 cells (Thermo) were cultured with Expi293 medium. The cells were routinely checked for mycoplasma contamination.

### Genes and Plasmids

The full-length coding sequence (CDS) of SARS-CoV-2 ORF8 (NC_045512.2) was prepared by gene synthesis (IDT). ORF8 CDS tagged with or without His was inserted into the pcDNA3.4 expression vector. Receptor The cDNA encoding IL-17RA (NM_00128905.2), IFNGR1 (NM_000416.3), IFNGR2 (NM_001329128.2) was cloned into a pME18S vector. All constructs were verified by DNA sequencing.

### Transfection

Plasmid DNA (0.8 μg in 50 μl of OPTI-MEM (Life Technologies) mixed with polyethylenimine “Max” (MW: 25,000) (Polysciences) (4 μg in 50 μl of OPTI-MEM) was incubated at room temperature for 20 min. This diluent was transfected into HEK293T cells at a density of 2 × 10^5^ cells/well in 24-well tissue culture plates.

### Establishment of Monoclonal Antibodies

Balb/c mice were immunized with human ORF8 protein and TiterMax Gold adjuvant. Two weeks after immunization, lymph node cells were fused with SP2/0, and hybridomas that recognized SARS-CoV2-ORF8-transfected HEK293T cells were obtained. Antibodies (#1-3-2, and #5-6-3) produced in culture supernatant were purified by protein A affinity chromatography.

### Detection of Soluble ORF8 by Flow Cytometry

Anti-ORF8 mAb (#1-3-2) was coupled to aldehyde/sulfate latex beads (Thermo Fisher Scientific A37304). The antibody was mixed with the latex beads and shaken at room temperature overnight. Then, the antibody-coupled beads were washed 3 times and suspended in PBS containing 0.1% glycine and 0.1% NaN3. Next, the antibody-coupled beads were mixed with the culture supernatants, virus supernatants, or patient serum for 30 mins. Subsequently, the beads were stained with biotinylated anti-ORF8 mAb (#5-6-3), and then these beads were stained with APC-labeled streptavidin (BD bioscience). A FACS Verse cytometer (Becton Dickinson) was used for analysis of the stained beads.

### PBMC and CD14 Monocyte Isolation and stimulation

The PBMCs derived from healthy donors were isolated from peripheral blood by Ficoll-Hypaque gradient separation. CD14-positive monocytes were isolated by using CD14 MicroBeads (Miltenyi). Cells were stimulated for different time periods with recombinant ORF8 protein, IFN-γ. For blockade of the IL-17 RA signaling, a monoclonal anti-human IL-17RA Ab (R004, Invitrogen) or irrelevant control anti-human MHC class II Ab (M5, Bio X Cell) was added to the cells 2h before stimulation.

### Recombinant proteins

The pcDNA3.4 expression vector containing the sequence that encodes the His-tagged extracellular domain of the ORF8 protein, and His-tagged human IL-17 and IFN-γ were transfected into Expi293 cells, then the His-tagged ORF8 protein produced in the culture supernatants was purified with a Talon resin (Clontech). The absence of endotoxin contamination was confirmed using the ToxinSensor chromogenic LAL Endotoxin Assay Kit (GeneScript).

### Assay for Cytokine Production

The amounts of cytokines in the supernatants were measured by a BD Human Inflammatory cytokine CBA kit (BD).

### Expression of His-tagged ORF8 and binding to cytokine receptors

HEK293T cells were transfected with full length of human IL-17RA, IFNGR (IFNGR1, IFNGR2), and IL-4R(IL-4RA, common-gamma chain) with or without His-tagged ORF8. Forty-eight hours after transfection, surface expression of ORF8 was detected by staining with anti-His mAb. For the detection of soluble ORF8 to cytokine receptors, His-tagged ORF8 was incubated with the receptor transfected cells (37°C for 1 hours), followed by anti-His antibody (37°C for 30 min), and samples were analyzed using FACSVerse (BD Bioscience).

### Viruses

SARS-CoV-2 was obtained from the Kanagawa Prefectural Institute of Public Health (KNG19-020). The stock virus was amplified in TMPRSS2-expressing VeroE6 cells. The SARS-CoV-2 infection assay was carried out in a Biosafety Level 3 laboratory.

### Generation of SARS-CoV-2 Recombinant Virus

SARS-CoV-2 recombinants were generated by CPER reaction as described previously (PMID: 33838744Ref;), with some modifications. In brief, a total of fourteen SARS-CoV-2 (2019-nCoV/Japan/TY/WK-521/2020) cDNA fragments (#1-#13) were amplified by PCR and subcloned into the pBlueScript KS(+) vector. The DNA fragment containing the CMV promoter, a 25-nt synthetic poly(A), hepatitis delta ribozyme and BGH termination, and polyadenylation sequences (#14) were synthesized from the Integrated DNA Technologies (Iowa, USA), and subcloned into the pBlueScript KS(+) vector. To generate a reporter SARS-CoV-2, we inserted a NanoLuc (NLuc) gene and 2A peptide into ORF6 of the viral genome of the #12 fragment (SARS-CoV-2/NLuc2AORF6). To generate an ORF8-deficient SARS-CoV-2(SARS-CoV-2/ΔORF8), the ORF8 gene was deleted by PCR. For the CPER reaction, fourteen DNA fragments that contained approximately 40- to 60-bp overlapping ends for two neighboring fragments were amplified by PCR from the subcloned plasmids. Then, the PCR fragments were mixed in equimolar amounts (0.1 pmoL each) and subjected to CPER reaction within 50 μL reaction volumes using PrimeSTAR GXL DNA polymerase (Takara Bio., Shiga, Japan) The cycling conditions were an initial 2 min of denaturation at 98°C; 35 cycles of 10 seconds at 98°C, 15 seconds at 55°C, and 15 min at 68°C; and a final extension for 15 min at 68°C. Half of the CPER product was transfected into IFNAR1-deficient HEK293 cells, in which human ACE2 and TMPRSS2 are induced by tetracycline (HEK293-3P6C33 cells) with TransIT-LT1 Transfection Reagent (Mirus, Madison, WI, USA), following the manufacturer’s protocols. At 24 h post-transfection, the culture supernatants were replaced with DMEM containing 2% FBS and doxycycline hydrochloride (1 μg/ml). At 7-10 days post-transfection, the culture supernatants containing progeny viruses (P0 virus) were passaged and amplified with VeroE6/TMPRSS2 cells.

### Animal Care and Use

All animal experiments with SARS-CoV-2 were performed in biosafety level 3 (ABSL3) facilities at the Research Institute for Microbial Diseases, Osaka University. The animal experiments and the study protocol were approved by the Institutional Committee of Laboratory Animal Experimentation of the Research Institute for Microbial Diseases, Osaka University (R02-08-0). All efforts were made during the study to minimize animal suffering and to reduce the number of animals used in the experiments. Four-week-old male Syrian hamsters were purchased from SLC (Shizuoka, Japan).

### SARS-CoV-2 ORF8 Administration to Syrian Hamsters

Recombinant ORF8 protein (100 or 200 μg), or control BSA (100 μg) were administered into the abdominal cavity of Syrian hamsters, and at 2 days post-administration all animals were euthanized, and blood and lungs were collected for histopathological examination and qRT-PCR.

### SARS-CoV-2 Infection into Syrian Hamsters

Syrian hamsters were anaesthetized with isoflurane and challenged with 6.0 × 10^4^ PFU via the intranasal route. Body weight was monitored daily for 5 days. At 5 days post-infection, all animals were euthanized, and lungs were collected for histopathological examinations and qRT-PCR.

### Quantitative RT-PCR

Total RNA of the lung homogenates was isolated using ISOGENE II (NIPPON GENE). Real-time RT-PCR was performed with the Power SYBR Green RNA-to-CT 1-Step Kit (Applied Biosystems) using a AriaMx Real-Time PCR system (Agilent). The relative quantitation of target mRNA levels was performed using the 2-ΔΔCT method. The values were normalized against those of the housekeeping gene, β-actin. The following primers were used: for β-actin; 5’-TTGCTGACAGGATGCAGAAG-3’ and 5’-GTACTTGCGCTCAGGAGGAG-3’, 2019-nCoV_N2; 5’- AAATTTTGGGGACCAGGAAC-3’ and 5’- TGGCAGCTGTGTAGGTCAAC -3’, IL-6; 5’-GGACAATGACTATGTGTTGTTAGAA- 3’and 5’- AGGCAAATTTCCCAATTG TATCCAG-3’, CCL3; 5’-GGTCCAAGAGTACGTCGCTG-3’and 5’- GAGTTGTGGAGGTGGCAAGG -3’, CCL5; 5’- TCAGCTTGGTTTGGGAGCAA -3’and 5’- TGAAGTGCTGGTTTCTTGGGT-3’, CXCL10; 5’- TACGTCGGCCTATGGCTACT -3’ and 5’- TTGGGGACTCTTGTCACTGG -3’.

### Hematoxylin and Eosin Staining

Lung tissues were fixed with 10% neutral buffered formalin and embedded in paraffin. Then, 2-μm tissue sections were prepared and stained with hematoxylin and eosin (H&E).

### Data and Statistical Analysis

FlowJo software (BD Bioscience) was used to analyze the flow cytometry data, and Graphpad Prism version 7.0e was used for graph generation and statistical analysis

## Results

### ORF8-knockout SARS-CoV-2 shows decreased pathogenesis

It is known that some viruses, such as poxvirus and herpesvirus, possess proteins that mimic cytokines to modulate immune response (15). The functions of the viral cytokines are diverse, and some of them are involved in immune evasion (15). It has been reported that the levels of proinflammatory cytokines and chemokines in COVID-19 patients infected with ORF8-deficient SARS-CoV-2 are lower than in those infected with wild-type SARS-CoV-2 containing ORF8 (16,17), and that ORF8 protein stimulation induced IL-6 production from monocyte cell line, THP-1 (9). These suggested that secreted ORF8 protein is involved in inflammation. To investigate the role of the ORF8 protein in the immune response after SARS-CoV-2 infection, we generated wild-type (WT) and ORF8-knockout SARS-CoV-2 viruses using the circular polymerase extension reaction (CPER) method (18-20) and compared their pathogenicity. SARS-CoV-2 replicates efficiently and causes pathological lesions in the lungs of Syrian hamsters similar to the features commonly reported in COVID-19 pneumonia patients (21-23). We analyzed the function of ORF8 protein by infecting hamsters with SARS-CoV-2 (Fig. 1A). There was no difference in viral titer in the lungs of hamsters infected with WT and ORF8-knockout SARS-CoV-2 (Fig. 1B). However, lymphocyte infiltration into the lungs and subsequent lung damage were less severe in the lungs of hamsters infected with the ORF8-knockout SARS-CoV-2 than in the lungs of those infected with WT SARS-CoV-2 (Fig. 1C). IL-6, CCL3, CCL5 and CXCL10 mRNA expression levels were significantly reduced in the lungs of hamsters infected with the ORF8-knockout SARS-CoV-2 (Fig. 1D).

**Fig.1.**
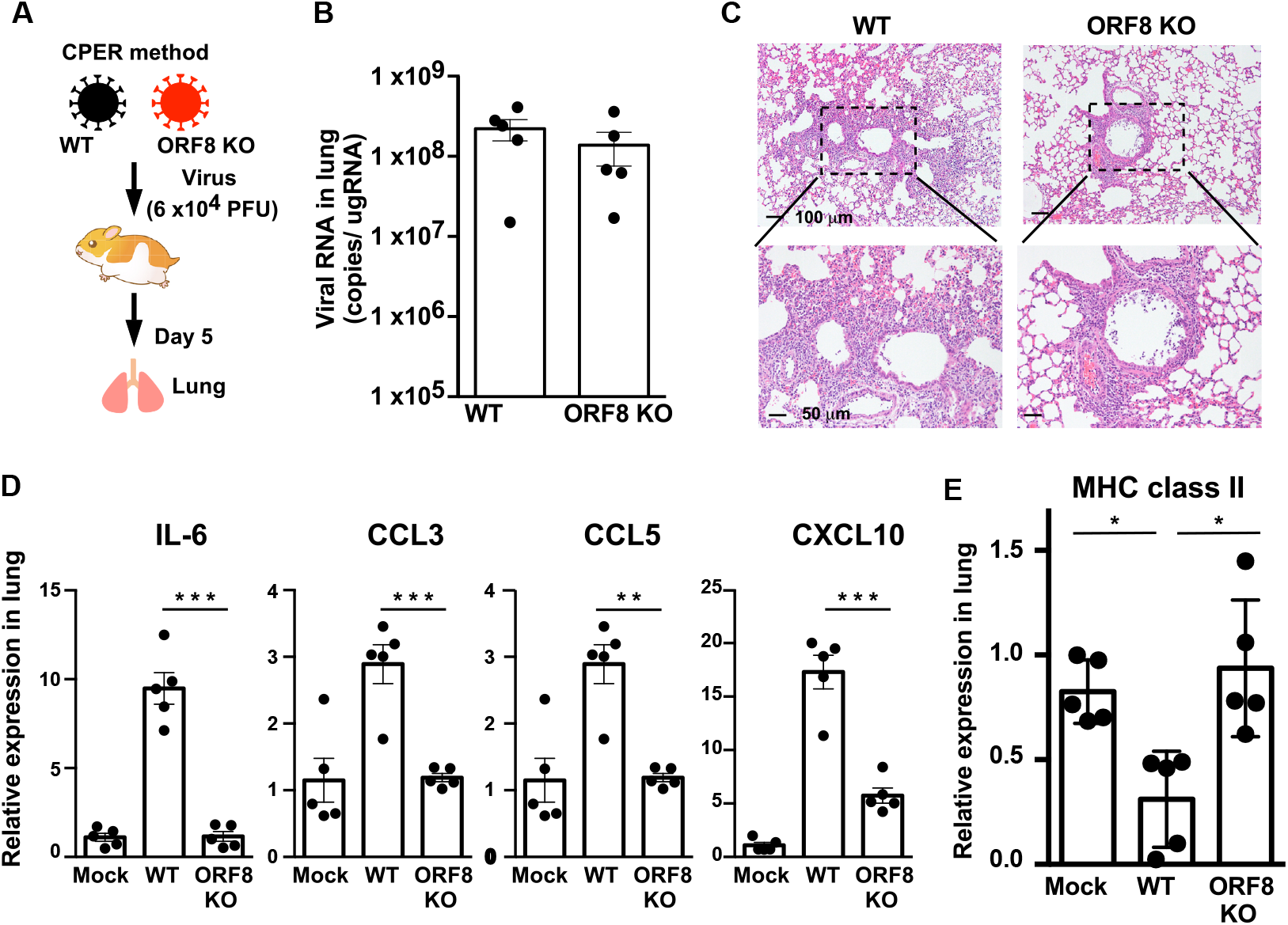
SARS-CoV-2 ORF8 regulates immune responses. (A) Schematic overview of the hamster infection experiment. (B) The ORF8-knockout SARS-CoV-2 (ORF8 KO) virus did not alter infectivity. Quantification of plaque-forming units (PFUs) from lung homogenates of infected hamsters at day 5 and genomic SARS-CoV-2 RNA as copies per μg of cellular transcripts. (C) H&E staining in the lung tissues of hamsters at 5 day post-infection. Scale bars, 100 μm. (upper panel) and 50 μm (lower panel). (D, E) Quantitative real-time PCR analysis of the transcript levels of inflammatory cytokine, chemokine (D) and MHC class II (E) in the lungs of infected hamsters at day 5. Results are representative of 5 mice/group from two independent experiments. Significance levels are shown as *p < 0.05, **p < 0.01, ***p < 0.005

The expression of major histocompatibility complex (MHC) class II molecules on monocytes is decreased in patients with severe COVID-19 compared to patients with mild disease (24-26). The MHC class II molecules play a critical role in antiviral adaptive immunity by presenting viral antigens to T cells (27). Virus escape from CD4^+^ T cell response is mediated by several mechanisms, with one of the mechanisms being the virus-mediated inhibition of the antigen presentation by MHC class II molecules (28). When we analyzed the expression of MHC class II on lung cells from on SARS-CoV-2-infected hamsters, MHC class II expression on lung cells from WT SARS-CoV-2-infected hamsters was down-regulated, but not that from ORF8-knockout SARS-CoV-2-infected hamsters (Fig. 1E). These results indicated that the ORF8 protein produced by SARS-CoV-2 infection is involved not only in the inflammatory response and subsequent lung damage, but also in the suppression of the acquired immune response via decreased MHC class II expression.

### SARS-CoV-2 ORF8 regulates immune responses

To confirm the role of the ORF8 protein in the immune response in vivo, recombinant ORF8 protein was administered to the hamsters and their lungs and PBMCs were then analyzed. Pulmonary lesions with inflammatory cell infiltration were observed in the lungs of ORF8 protein-administered hamsters, but not in PBS- or BSA-administered hamsters (Fig. 2A). In the lungs and PBMCs of hamsters administered the recombinant ORF8 protein, the transcription levels of IL-6, CXCL10, and CCL5 were up-regulated. CCL3 mRNA expression was also observed, although not at significant levels (Fig. 2B). Furthermore, we observed a decrease in the expression of the MHC class II gene in the lungs of hamsters treated with ORF8 protein compared to that in hamsters treated with control PBS or BSA (Fig. 2C). These results directly indicated that ORF8 protein is involved in dysregulation of immune responses.

**Fig. 2.**
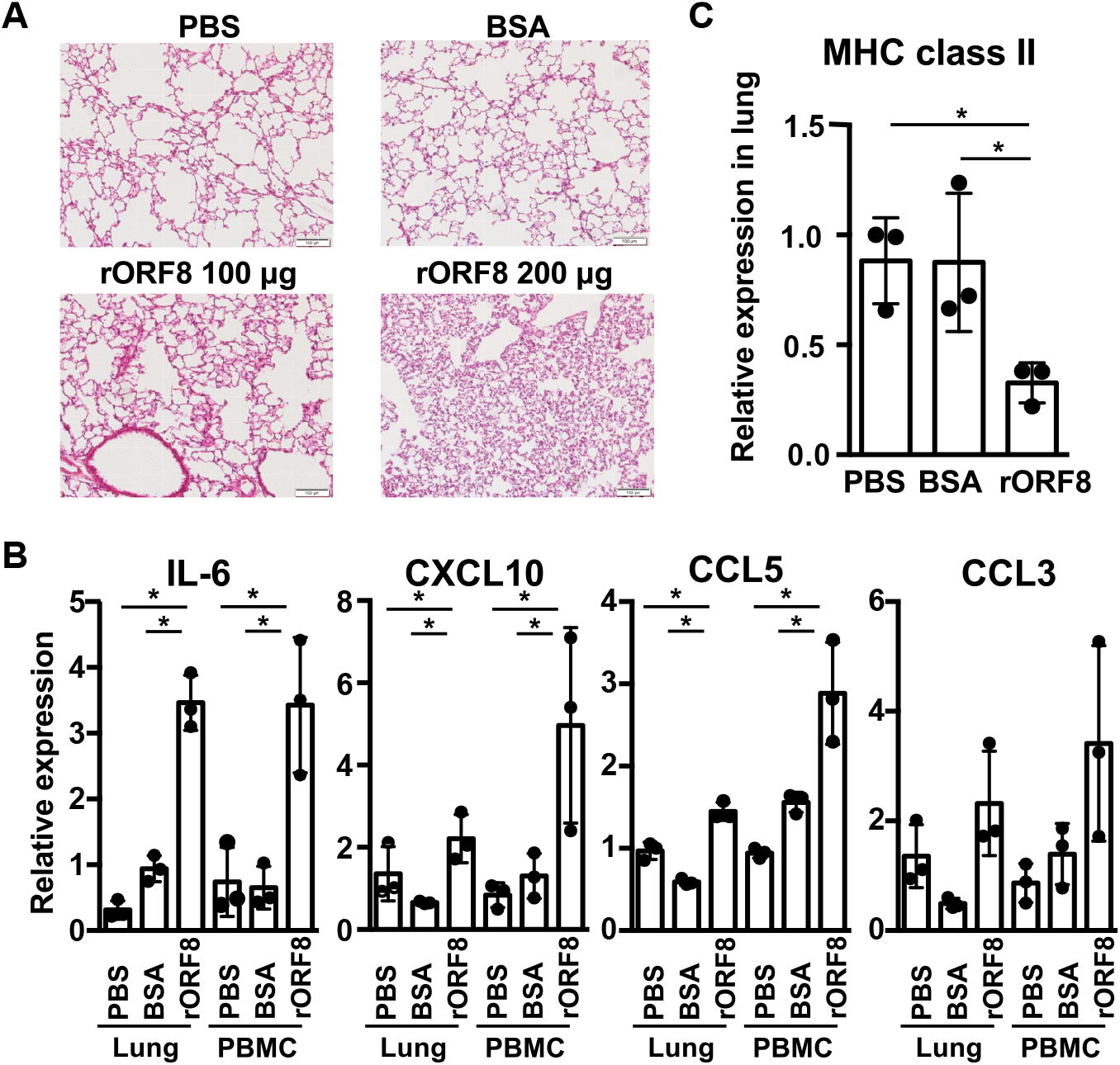
Recombinant ORF8 protein induced inflammation and lung tissue damage in hamsters. Recombinant ORF8 protein (rORF8, 100 or 200 μg), control BSA (100 μg), or PBS was administered into the peritoneal cavity of hamsters. Representative H&E staining of hamster lung tissue 2 days post-administration. Scale bars, 100 μm. (B, C) Quantitative real-time PCR analysis of mRNA expression of inflammatory cytokines and chemokines (B), and MHC class II (C) from lung tissue or PBMCs at day 2. Results are representative from two independent experiments. Significance levels are shown as *p < 0.05.

When monocytes from the PBMCs of healthy donors were stimulated with recombinant ORF8 protein, inflammatory cytokine, particularly high levels of IL-6 and IL-8, production was also observed (Fig. 3A). It has been reported that SARS-CoV2 ORF8 physically interacts with IL-17 receptor A (IL-17RA) (9,29,30). Consistent with these previous reports, IL-6 and IL-8 production induced by ORF8 protein were reduced with anti-IL-17RA antibody treatment (Fig. 3B). IFN-γ regulates the expression of MHC class II molecules on almost all cells including monocytes (31). We examined whether ORF8 protein could bind to IFN-γ receptor (IFNGR). IL-17RA, IFNGR, and IL-4 receptor (IL-4R) were expressed on HEK293T (Fig.S1), and examined the binding of ORF8 protein to these receptors. We found that in addition to IL-17RA, ORF8 protein bound to IFNGR but not to IL-4R (Fig. 3C). Next, we analyzed the effect of ORF8 protein on IFN-γ-induced MHC class II expression. MHC class II expression on THP-1 cells was enhanced by IFN-γ treatment, and this enhancement was attenuated by treatment with ORF8 protein (Fig. 3D). These results suggested that the ORF8 protein is involved in down-regulation of MHC class II through the modulation of IFN-γ signaling.

**Fig. 3.**
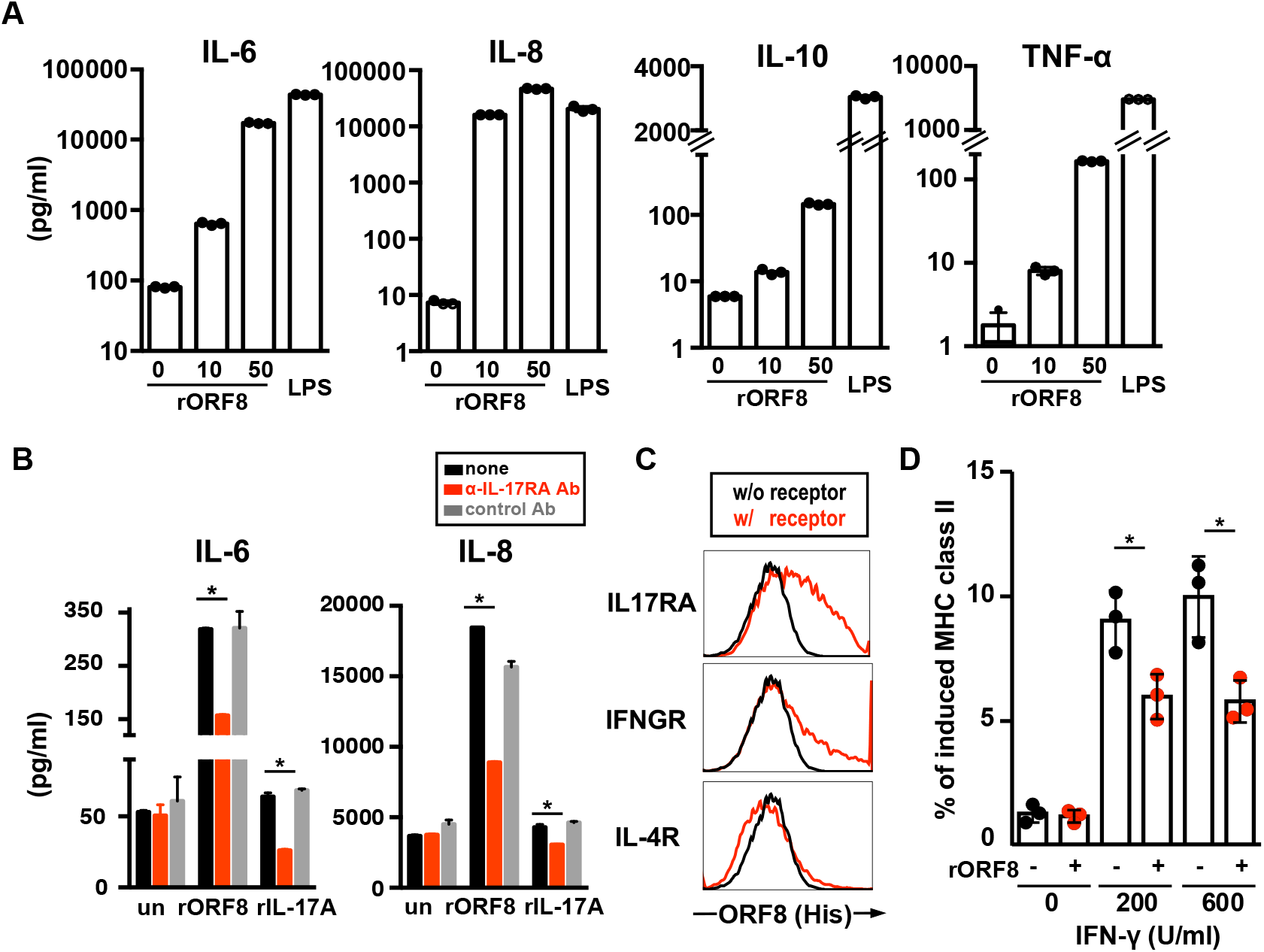
Recombinant ORF8 protein induced an inflammatory response and down-regulated MHC class II expression on human monocyte. (A) Recombinant ORF8 induced inflammatory cytokine production from human monocyte. Monocytes isolated from healthy donor PBMCs were stimulated with ORF8 protein (10, 50 μg/ml) or LPS (100 ng/ml), and cytokines in 24-hour-cultured supernatants were measured. The experiments were performed at least three times, and representative results from one experiment are shown. Error bars indicate SD of technical triplicates. (B) IL-6 and IL-8 production were through IL-17RA. Monocytes isolated from healthy donor PBMCs were stimulated with ORF8 protein (10 μg/ml) in the presence of anti-IL-17RA or control antibodies, and IL-6 and IL-8 in 24-hour-cultured supernatants were measured. The experiments were performed at least twice, and representative results from one experiment are shown. Error bars indicate SD of technical triplicates. Significance levels are shown as *p < 0.05. (C) MHC class II expression induced by IFN-γ was attenuated in ORF8 protein pre-treated THP-1cells. THP-1 was incubated with recombinant ORF8 (20 µg/ml). At 24 hours after, cells were stimulated with IFN-γ for 24 hours, and the expression of MHC class II was measured by flow cytometry. The experiments were performed three times, and representative results from one experiment are shown. Error bars indicate SD of technical triplicates. Significance levels are shown as *p < 0.05.

### SARS-CoV-2 ORF8 is a secretory protein detected in the body fluids of COVID-19 patients

To understand the role of ORF8 protein in COVID-19 patients, we examined the levels of ORF8 protein produced in COVID-19 patients. We generated monoclonal antibodies recognizing the ORF8 protein and established a highly sensitive ORF8 protein detection system (Fig. 4A). The spike protein was not detected by this ORF8 detection system at all (Fig. 4B), and the sensitivity of this system is quite high to detect ORF8 even at concentrations as low as 1 pg/ml (Fig. 4C). We then examined whether ORF8 proteins were secreted from SARS-CoV-2-infected cells. HEK293 cells expressing doxycycline (DOX)-induced ACE2 (18) were infected with SARS-CoV-2, and ORF8 protein secretion was examined. High levels of ORF8 protein (105 ng/ml) were detected in the culture supernatant of SARS-CoV-2-infected cells, but not from non-ACE2-expressing cells (Fig. 4D). Similar to previous findings of the SARS-CoV-2 ORF8 transfectant experiment (14), the ORF8 protein was detected in the culture supernatant at 6 hours post-infection, and the amount of ORF8 protein increased over time. The ORF8 protein was detected in the culture supernatant of WT SARS-CoV-2 infected cells but not in the culture supernatant of ORF8-knockout SARS-CoV-2-infected cells (Fig. 4E)

**Fig. 4.**
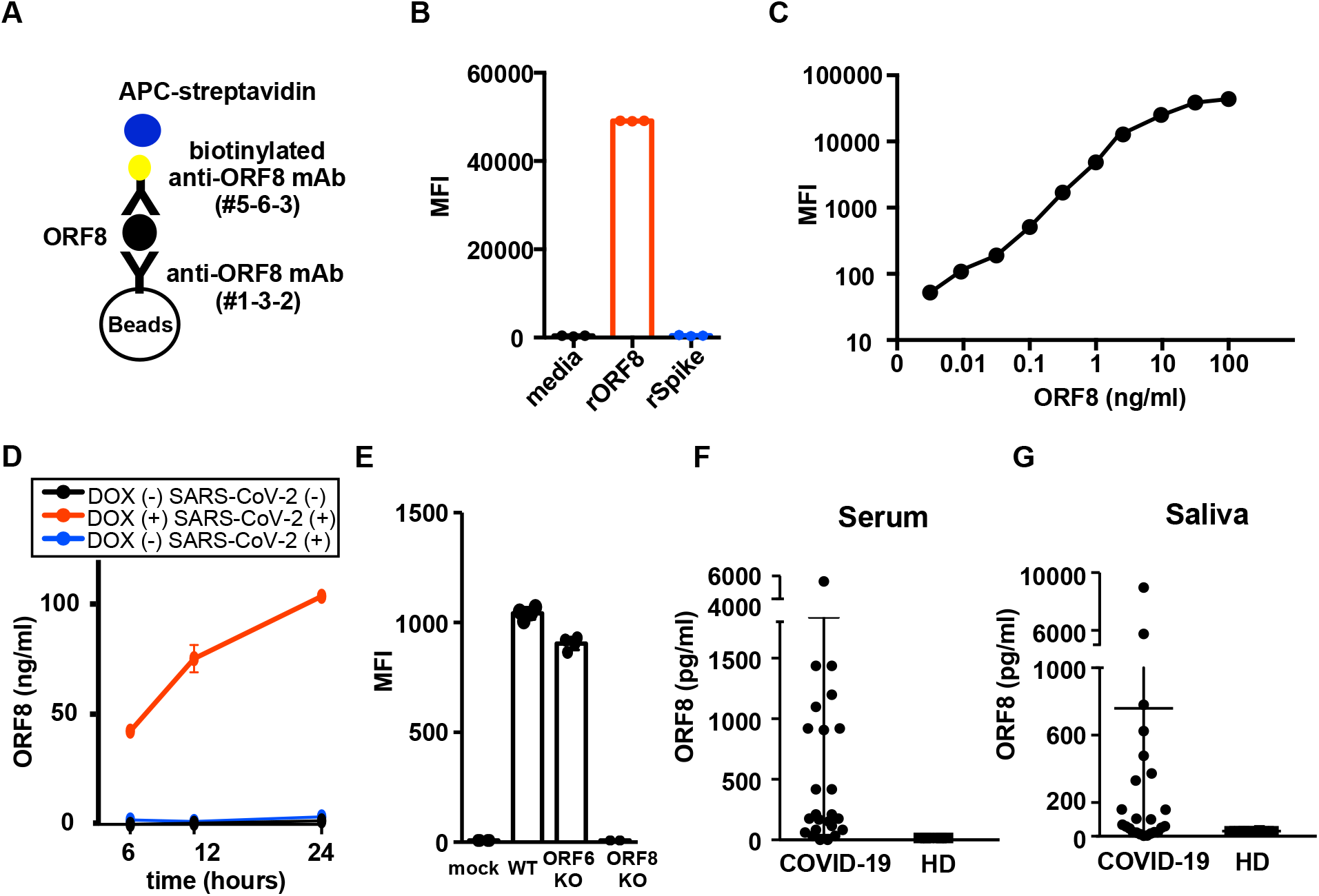
SARS-CoV-2 ORF8 is a secretory protein detected in serum and saliva of COVID-19 patients. (A) Schematic diagram illustrating the system for soluble ORF8 protein detection by anti-ORF8 mAb-coated beads and biotinylated anti-ORF8 mAb followed by streptavidin-APC. The ORF8 protein, but not Spike protein, was recognized by the anti-ORF8 mAb-bead assay. The recognition of recombinant ORF8 (100 ng/ml) and Spike proteins (100 ng/ml). (C) Standard curve generated using the anti-ORF8 mAb-bead assay; ORF8 concentration ranging from 1 pg/ml to 100 ng/ml. (D) the ORF8 protein was detected in the culture supernatant of SARS-CoV-2 virus-infected cells. ACE2-induced (Dox (+)) or not induced (Dox (-)) HEK293 cells were infected with SARS-CoV-2 virus and the ORF8 protein secreted in the culture supernatants was analyzed by flow cytometry. Results are representative of 2 independent experiments. (E) The ORF8 protein is deleted in ORF8-knockout SARS-CoV-2 (ORF8 KO) virus. The amount of ORF8 protein in the culture supernatants of ORF8 KO-, or control wild-type- and ORF6 KO-virus infected ACE2 expressing HEK293 was analyzed by flow cytometry. Results are representative of 2 independent experiments. (F, G) SARS-CoV-2 ORF8 was detected in the sera and saliva of COVID-19 patients. The amount of SARS-CoV2-ORF8 in the serum (F) and saliva (G) of hospitalized patients (n=24) and healthy control donors (n=10) was examined by flow cytometry.

As the ORF8 protein was detected at high levels in the culture supernatant of SARS-CoV-2-infected cells in our system, we addressed whether the ORF8 protein could be detected in the body fluids from COVID-19 patients. Consistent with previous report, the ORF8 protein was detected in the serum of COVID-19 patients, as the serum from 22 of 24 (92%). Moreover, ORF8 protein was detected in the saliva from 17 of 24 (70%) COVID-19 patients (Fig. 4F and 4G). The levels of ORF8 varied among individuals, with some patients producing as little as 30 pg/ml and others producing more than 6000 pg/ml. We then evaluated the kinetics of the serum levels of ORF8 protein as well as IL-6 levels and anti-ORF8 antibody titers in five COVID-19 patients. The ORF8 protein was detected relatively early in the course of infection, and the levels of IL-6 were found to be elevated immediately after ORF8 protein detection. Anti-ORF8 antibody was increased at about one week after ORF8 protein was detected (Fig. 5A). We examined the levels of LDH, CRP, and ferritin, which are also biomarkers of inflammation (32-34), and found that the peak levels of these biomarkers tended to come after the peak in ORF8 production, similar to that observed for IL-6 production (Fig. 5B). These results suggested that the ORF8 protein is a secretory protein involved in inflammation.

**Fig. 5.**
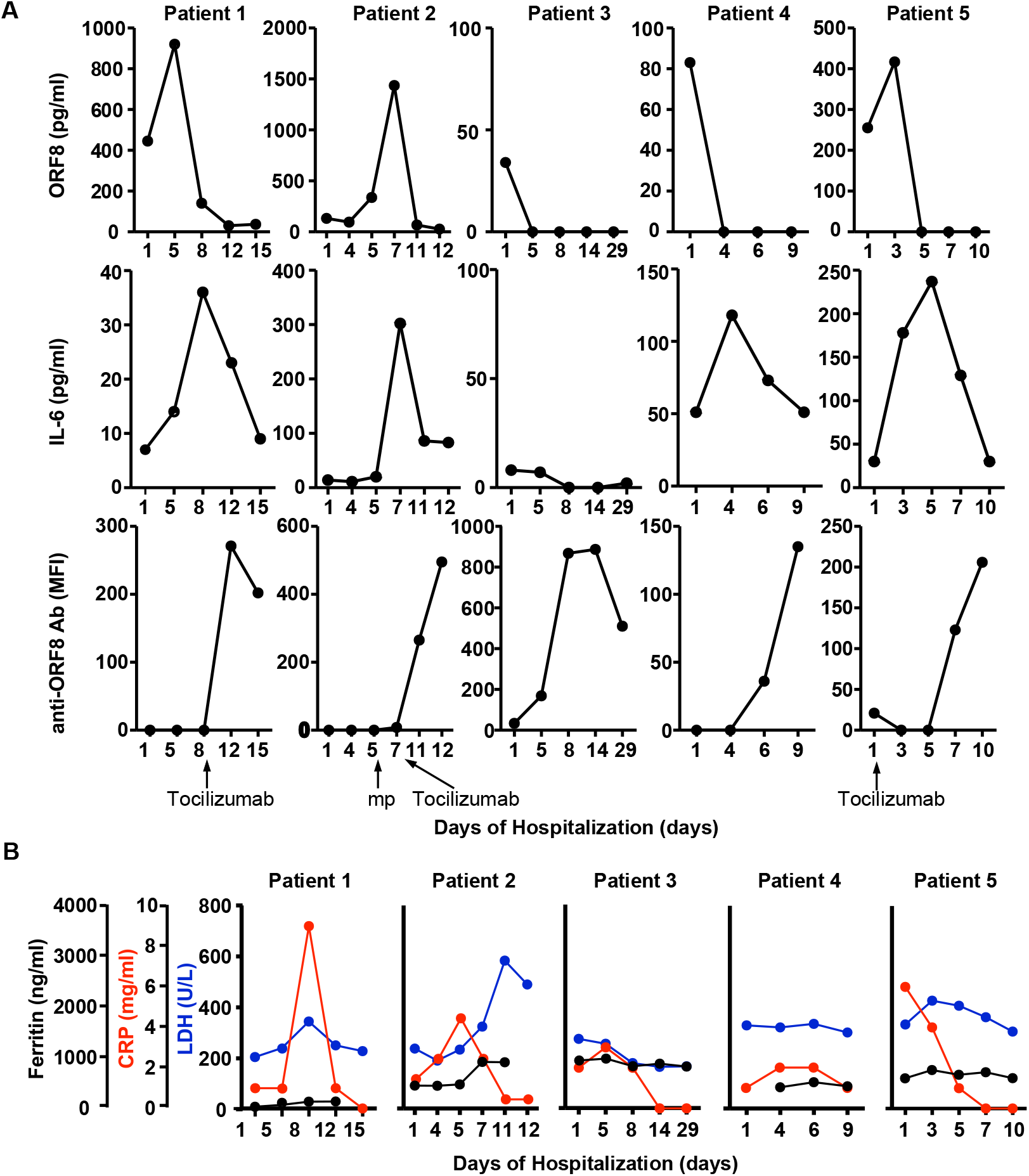
Correlation between ORF8 protein and biomarkers of inflammation in COVID-19 patient serum. (A) Changes in ORF8 protein, IL-6, and ORF8-specific antibody in the sera. The amount of ORF8 protein, IL-6, and anti-ORF8 antibody collected longitudinally from COVID-19 patients. Patients who received tocilizumab or methylprednisolone sodium succinate (mp) treatment are shown in the figure. (B) Changes in serum LDH (blue), CRP (red), and ferritin (black) levels in the sera collected longitudinally from COVID-19 patients. CRP, C-reactive protein; LDH, lactate dehydrogenase.821)

## Discussion

Inflammation is a key feature of COVID-19, and excessive inflammation leads to unfavorable outcomes and even death. Therefore, it is quite important to identify the factors causing the excessive inflammatory response in SARS-CoV-2 infection. In the present study, we analyzed the physiological function of ORF8 protein by generating ORF8-knockout SARS-CoV-2. We found that the lung inflammation observed in wild-type SARS-CoV-2-infected hamsters was decreased in ORF8-knockout SARS-CoV-2-infected hamsters. Similar results were obtained by administration of recombinant ORF8 protein to hamsters.

MHC class II molecules play an important role in both cellular and humoral immune responses by presenting peptide antigens to helper T cells. However, the expression levels of MHC class II molecules have been reported to decrease in severe COVID-19 patients (24-26), suggesting that certain factors in SARS-CoV-2 are involved in MHC class II down-regulation. Consistent with the observation in COVID-19 patients, we found that MHC class II expression is down-regulated in the lungs of WT but not ORF8-deficient SARS-CoV-2-infected hamsters as well as in recombinant ORF8-administered mice. These data indicated that ORF8 is involved in the immune evasion of SARS-CoV-2 by the down-regulation of MHC class II expression. It is noteworthy that the SARS-CoV-2 ORF8 protein allows the host to protect themselves from the virus by inducing an inflammatory response, while the virus attempts to escape the host’s immune system by suppressing the expression of MHC class II molecules.

ORF8 protein-induced IL-6 and IL-8 production were attenuated by anti-IL-17RA antibody treatment, but not completely. ORF8 protein has been reported to bind to several other cytokine receptors (29,35), and we found that ORF8 protein could bind to IFNGR. These results suggested that ORF8 protein might bind to other unknown receptors as well as the IL-17RA to induce inflammatory cytokine, or soluble ORF8 protein might be directly incorporated into the cells and function intracellularly. Therefore, more work will be needed to understand how ORF8 protein regulates immune responses in the future, and these analysis of the ORF8 protein is important to understand the pathogenicity of SARS-CoV-2.

Here, we also found that serum levels of IL-6, CRP, LDH, and ferritin, which are biomarkers of inflammation in COVID-19 patients (32-34), were well correlated with the ORF8 protein levels. It has been reported that SARS-CoV-2 infects not only respiratory organs but also various tissues such as kidney, small intestine, pancreas, and brain (36-38). Infection to non-respiratory organs is difficult to detect using current detection system such as PCR. However, ORF8 protein will be secreted from any SARS-CoV-2 infected cells. Therefore, ORF8 protein would be a useful clinical marker to detect SARS-CoV-2 infection in non-respiratory organs, although further large scale clinical studies are required to validate the usefulness to measure ORF8 levels. Our results indicate that monitoring serum ORF8 protein levels in COVID-19 patients is important in the evaluation of disease condition. Also, inhibitors of the ORF8 protein might be useful in blocking SARS-CoV-2-induced inflammation. Further studies on the ORF8 protein would contribute to a better understanding of the etiology of COVID-19.

## Author contribution

M.K., E.M., T.O., and H.A. conceived experiments. M.K., S.K., W.N., S.M., K.I.,, Y.L., T.S., E.M., and T.O. performed experiments. A.N., N.A., N.H., S.O., K.T, and K.O. collected patients’ sera and Saliva. M.K., T.O., and H.A. wrote the manuscript. All authors read, edited, and approved the manuscript.

